# Molecular and network disruptions in neurodevelopment uncovered by single cell transcriptomics analysis of *CHD8* heterozygous cerebral organoids

**DOI:** 10.1101/2023.09.27.559752

**Authors:** Maider Astorkia, Yang Liu, Erika M. Pedrosa, Herbert M. Lachman, Deyou Zheng

**Author notes:** Correspondence should be addressed to, Deyou Zheng, Ph.D., Albert Einstein College of Medicine, 1301 Morris Park Ave. Bronx, NY 10461, Herbert Lachman, M.D., Albert Einstein College of Medicine, 1300 Morris Park Ave. Bronx, NY 10461. equal contribution (Maider Astorkia, Yang Liu). Division of Allergy, Pulmonary, and Critical Care Medicine, Department of Medicine and Department of Biostatistics, Vanderbilt University Medical Center, Nashville, TN, USA.

## Abstract

About 100 genes have been associated with significantly increased risks of autism spectrum disorders (ASD) with an estimate of ∼1000 genes that may be involved. The new challenge now is to investigate the molecular and cellular functions of these genes during neural and brain development, and then even more challenging, to link the altered molecular and cellular phenotypes to the ASD clinical manifestations. In this study, we use single cell RNA-seq analysis to study one of the top risk gene, *CHD8*, in cerebral organoids, which models early neural development. We identify 21 cell clusters in the organoid samples, representing non-neuronal cells, neural progenitors, and early differentiating neurons at the start of neural cell fate commitment. Comparisons of the cells with one copy of the *CHD8* knockout and their isogenic controls uncover thousands of differentially expressed genes, which are enriched with function related to neural and brain development, with genes and pathways previously implicated in ASD, but surprisingly not for Schizophrenia and intellectual disability risk genes. The comparisons also find cell composition changes, indicating potential altered neural differential trajectories upon *CHD8* reduction. Moreover, we find that cell-cell communications are affected in the *CHD8* knockout organoids, including the interactions between neural and glial cells. Taken together, our results provide new data for understanding CHD8 functions in the early stages of neural lineage development and interaction.

## Introduction

*CHD8* (**c**hromodomain **h**elicase **D**NA binding protein 8) is a ubiquitously expressed member of the CHD family of ATP-dependent chromatin-remodeling factors that play important roles in chromatin dynamics and transcription. It is one of the strongest risk genes whose loss-of-function mutations have been associated with Autism Spectrum Disorder (ASD), a class of neurodevelopmental disorders characterized by persistent deficits in social communication/interaction and stereotypical behaviors/interests (DSM-5). It was initially discovered from studies of rare *de novo* mutations present in ASD probands but absent in their healthy siblings or parents, using whole exome or whole genome sequencing ^1–4^. The strong association has since been reproduced by many subsequent ASD genetics studies using large cohorts consisting of thousands of samples ^5–8^. Some studies further suggest that ASD individuals with *CHD8* mutations may define a subgroup of ASD, with stereotypical ASD features, early brain outgrowth, and gastrointestinal syndromes^9,10^.

Many studies have been carried out to understand CHD8 functions in neural and brain development, with additional work in non-neural contexts (e.g., cancers), using human cell, mouse, rat, or zebrafish models ^9,11,20–26,12–19^. These studies found that CHD8 bound to thousands of genes, largely biased to gene promoters, and reduced CHD8 expression resulted in disruptions of gene networks involved in neurodevelopment, such as WNT signaling. A meta-analysis further defined the conserved roles of CHD8 across human, mouse, rat, *in vivo* and *in vitro* models. It showed that CHD8 reduction affected a large number of genes involved in fundamental biological processes (e.g., neuronal development and function, cell cycle, chromatin dynamics, and metabolism) ^27^. The analysis also found that differentially expressed genes in the compared studies, however, were distinct and often not overlapping, indicating that genes targeted by CHD8 are likely context dependent, like other chromatin modifiers. Consistent with this, the reported molecular, cellular and behavior phenotypes in multiple *Chd8* mouse models also varied, with mostly small changes ^22^. One caveat in the transcriptomics analyses in those studies is the usage of bulk tissues, which may have masked cell type specific effects.

With an advantage in studying the role of CHD8 in multiple human cell types simultaneously, several research groups, including us, have used cerebral organoids to model and decipher CHD8 function in early human neural and brain development ^14,28,29^. Coupling its usage with single cell RNA sequencing (scRNA-seq) ^28–30^, investigators have uncovered both common and cell type specific genes that were affected by *CHD8* haploinsufficiency, generated, for example, using CRISPR editing to knock out one copy of *CHD8* (*CHD8*^+/-^) or introduce patient specific mutations. While the studies consistently point to a role of CHD8 in regulating neural development, the cell types and neural differentiation stages altered in CHD8-disruption lines seem to vary. Moreover, differences among cell lines in the same study were also noticeable. Such variations, although not very satisfactory, are quite commonly observed in brain organoid studies, partially due to the sensitivity of the neural differentiation system to genetic background, differentiation protocols, and stochasticity.

Considering the variations in molecular and cellular phenotypes that were observed in previous CHD8 studies of both mouse models and human organoid models, we decided to perform an independent scRNA-seq analysis of the cerebral organoids that we generated from induced pluripotent stem cell lines (iPSCs) of a typically developing individual and the corresponding isogenic lines with one copy of *CHD8* knocked out using CRISPR-Cas9 gene-editing technology. The generation of the lines and bulk RNA-seq analysis of the cerebral organoids and 2D neural samples were described in our previous reports, which showed that genes highly expressed in inhibitory neurons were affected by CHD8 haploinsufficiency ^13,14^. Here, we characterize the cell types in the organoids, neural differential trajectory and cell-cell communication, and their alterations upon CHD8 reduction. Distinct from studies conducted previously on cerebral organoids from longer culture periods ^28–30^, current work analyzes organoids after only 50 days of neural differentiation, thus providing complementary and new insights into the role of CHD8 in early neural differentiation.

## Material and Methods

### Cerebral organoid preparation

Self-organizing cerebral organoids were derived from induced pluripotent stem cells (iPSC) using the STEMdiff™ Cerebral Organoid Kit developed by STEMCELL Technologies™ (catalogue # 08570), which is based on the protocol developed by Lancaster et al. ^31^. The protocol uses defined, serum-free cell culture media. The first step (days 0-5) is the development of embryoid bodies (EBs). iPSCs were grown in a 6-well plate at 70 - 80% confluence. Areas of spontaneous differentiation were manually removed by scraping with a pipette tip. Medium was aspirated and the iPSC cells were washed with 1 mL of sterile phosphate-buffered saline (PBS). Following aspiration of PBS, 1 mL of Gentle Cell Dissociation Reagent was added and the cells were incubated at 37°C for 8 - 10 minutes. Using a 1 mL pipettor, the cells were gently resuspended by pipetting up and down slowly 3 - 5 times. The cells were transferred to a sterile 50 mL conical tube. The well was rinsed with an additional 1 mL of EB Seeding Medium (EB Formation Medium + 10μM Y-27632) and added to the tube containing cells. Following centrifugation of cells at 300 x g for 5 minutes, supernatant was removed and discarded, and 1 - 2 mL of EB Seeding Medium was used to resuspend cells. The cells were counted in a hemocytometer using Trypan Blue staining. The volume of cells required to obtain 90,000 cells/mL was determined; this was added to an appropriate volume of EB Seeding Medium. Using a multi-channel pipettor,100 μL of the cell suspension was added to each well of a 96-well round-bottom, ultra-low attachment plate (9000 cells/well). The cells were then incubated at 37°C for 24 hours, undisturbed, in a humidified incubator containing 5% CO_2_. Under an inverted microscope, small EBs (∼100 - 200 μm) were visible. On days 2 and 4, 100 μL of EB Formation Medium was gently added to each well using a multi-channel pipettor. On day 5, the EBs reached a diameter of 400 - 600 μm. On day 5, induction medium at room temperature was prepared according to the manufacturer, and 0.5 mL was added to each well of a 24-well ultra-low attachment plate. 1 - 2 EBs were added to each well using a wide-bore 200 μL pipette tip, drawing up 50 μL from one well of the 96-well plate. Most of the medium was carefully removed by ejecting it back into the well, retaining EBs in the pipette tip. The EBs were dispensed into one well of the 24-well plate containing Induction Medium. The plates were then incubated at 37°C for 48 hours. On day 7, Matrigel® was thawed on ice for 1 - 2 hours and used at 1.44 mL Matrigel® (15 μL/well x 96 wells).

Expansion Medium was prepared according to the manufacturer’s instructions and warmed to room temperature. A UV-sterilized embedding surface (Parafilm®) was placed into an empty, sterile, 100 mm dish. EBs were drawn up using a wide-bore 200 μL pipette tip, along with 25 - 50 μL of medium, and transferred to the embedding surface. This was repeated until 12 - 16 EBs were collected on the embedding surface. Excess medium was removed from each EB by drawing up medium with a standard 200 μL pipette tip. 15 μL of Matrigel® drawn up in a cold 200 μL standard pipette tip was added drop by drop onto each EB. The EBs were repositioned to the center of the droplet using a new, cold 200 μL pipette tip, The plate was then placed in an incubator at 37°C for 30 minutes to polymerize the Matrigel®. Sterile forceps were used to grasp the embedding surface containing Matrigel® droplets. The sheet was positioned directly above one well of a 6-well ultra-low adherent plate. Using a 1 mL pipettor, Expansion Medium was drawn up and the Matrigel® droplets were gently washed off the sheet and into the well. Using 3 mL of Expansion Medium/well, the process was repeated until all 12 - 16 Matrigel® droplets were in the well. The plates were then incubated at 37°C for 3 days, after which embedded organoids were observed to develop expanded neuroepithelia (budding of the EB surface).

From days 10-40, the organoids were grown in Maturation Medium, which was prepared according to the manufacturer’s instructions. Expansion Medium was slowly removed with a 5-10 mL serological pipette. This was replaced with Maturation medium (3mL/well). Maturation Medium was kept at room temperature prior to its addition to the organoids. The plate of organoids was placed on an orbital shaker in a 37°C incubator. A full medium change was made every 72 hours.

From day 40 to day 50, the organoids were cultured with STEMdiff™ Cerebral Organoid Maturation Kit (Catalog #08571) with a full medium change every 48 hours. The organoids were harvested for single cell RNA sequencing on day 50.

### Single cell RNA-seq data acquisition

Single cell RNA-seq libraries were prepared with a 10x Chromium (10x Genomics, Pleasanton, CA) using the Chromium Next GEM Single Cell 3’ GEM, Library & Gel Bead Kit v3.1 (10x Genomics, Cat# PN-1000121), according to the manufacturer’s instructions, at the Einstein Genomics Core Facility, followed by paired-end sequencing on an Illumina NextSeq 500 platform. The four barcoded samples were pooled and sequenced together.

### Single cell RNA-seq data analysis

Cell Ranger software (v3.0.2; 10X Genomics) ^32^ with default parameters was used to align the scRNA-seq reads to the human GRCh38 reference genome, identify cell barcodes, and generate gene expression count matrices. The count matrices were analyzed by the RISC (v1.0) ^33^ package for each of the four samples and their integration. Cells with a minimum of 1000 and a maximum of 40000 UMI (unique molecular identifier) counts, as well as a minimum of 200 genes were kept for further analysis. Data integration was performed by the RPCI (Reference Principal Component Integration) method ^33^, using the iPS1 sample as the reference and the top 20 principal components (PCs). Dimensionality reduction and cell clustering were performed on the integrated data by UMAP (Uniform Manifold Approximation and Projection) and Louvain methods, respectively, with the top 20 PCs, using RISC (default parameters). One small cell cluster very distant from other cells was identified as endothelial cells by marker genes (e.g., *PECAM1*) and removed from further analysis. Gene markers for each cluster were identified using *AllMarker* function and Quasi-Poisson method. Differential expression analysis between *CHD8* KO and controls was run by comparing the cells from the two KO samples with cells from the two iPS control samples. Differentially expressed genes (DEGs) were determined using the *scDEG* function (using Negative Binomial option) at log2(fold change) >1 and adjusted p-value (padj) < 0.05. Difference in cell population abundance between *CHD8* KO and controls was determined by miloR (v1.1.0) ^34^ using 30 k-nearest-neighbors (KNN) for graph building and 30 dimensions for KNN (K nearest neighbors) refinement.

### Pseudotime trajectory analysis

Spliced/unspliced count matrices for each of the samples were generated by velocyto ^35^, using the GRCh38 as the reference and repeat masked. The resulting loom files were further processed by scVelo (v0.2.4) ^36^ to compute and visualize the proportions of spliced/unspliced reads for each of the cell clusters within each sample. Variable genes were detected by minimum number of counts and dispersion, and data across cells were normalized by total library sizes and logarithm-transformed using default parameters through *pp*.*filter_and_normalize* function. Mean and variance of the data were calculated by nearest neighbors in PCA space using 30 PCs and 30 neighbors, and the graph was embedded into two dimensions UMAP. Clustering of the data was based on Louvain statistics. The estimation of RNA velocity was obtained by *tl*.*velocity* and *tl*.*velocity_graph* functions using default parameters, while projection of RNA velocity to the UMAP was obtained from a dynamic function (*tl*.*umap*) together with the RISC clustering. The *tl*.*louvain* function was also used to find clusters based on dynamic data to help cell type identification. Finally, the initial and terminal states were identified by CellRank (v1.5.1) ^37^ using *tl*.*terminal_states* and *tl*.*initial_states*.

### Cell type annotation

Cell type identification was based on a combination of known markers, marker enrichment and pseudotime results. First, neuronal and non-neuronal cell types were separated by the expression of *GAP43, STMN2, DCX* (neuronal) and *VIM, HES1, SOX2* (non-neuronal), as described by Tanaka et al. ^38^. Neuronal clusters were then classified as excitatory cortical neurons through *SLC17A7, TBR1* and *NEUROD2* expression and inhibitory interneurons by *SLC32A1, GAD1* and *GAD2* markers. Neuronal progenitors were identified by high expression of corresponding marker genes and their positions in the pseudotime trajectory. Cells highly expressing *TOP2A* and *MKI67* were annotated as neuroepithelial cells, while astrocytes were identified by high expression of *GFAP* and *SLC1A3*. Astrocytes progenitors were detected through lower expression of Astrocytes markers using RNA velocity as support. In the same way, oligodendrocyte cells and their progenitors were identified by high and low *OLIG1* expression, respectively. CRYM cluster was identified by unique *CRYM* expression and UPRC cells by *DDIT3* expression and additionally enrichment of its markers in GO:0006986 term (“response to unfolded protein”). Epithelial cells were identified by markers and the enrichment of “NS-moderated 16-33-Epithelial-Ciliated term” on ToppCell Atlas ^39^. Lastly, mesenchymal cells were identified by *PIFO* and *PCP4* expression and enrichment of markers differentiating mesenchymal cilium from mesenchymal BMP cells.

### Over representation analysis based on curated gene sets and Gene Ontology

Two over representation analysis were performed for DEGs between *CHD8* KO and controls, for each of the cell types independently. In the first, *EnrichGO* function from clusterProfiler ^40^ was used to identify overrepresented GO terms in *“biological processes,”* with *simplify* function used to reduce redundancy. In the second, we tested the overrepresentation of our DEGs in gene sets associated with ASD, schizophrenia (SCZ), intellectual Disability (ID), Attention Deficit Hyperactivity Disorder (ADHD), and neurodevelopmental disorders, for a total of seven gene sets curated in our previous work ^41^. Fisher test was applied using GeneOverlap ^42^ package.

### Pathway enrichment analysis

In addition to overrepresentation analysis, DEGs were also ranked by their expression fold changes and used for Gene Set Enrichment Analysis (GSEA, v4.2.2) ^43,44^. Gene sets in the GO:*”biological processes”* were selected for enrichment analysis at FDR < 25 % (default).

### Cell-Cell interaction

CellChat ^45^ software was used to infer ligand-receptor (L-R) interactions between the identified cell types in control and KO samples. For that, normalized gene expression matrix from the RISC integration was loaded to CellChat, with iPSC1 and iPSC2 data merged to one control group while KO1 and KO2 to one KO group, and then *computeCommunProb* function was used to select genes expressed in > 20% of cells within one cell type for determining statistically significant (p < 0.05, permutation test) L-R interactions. From those L-R interactions, CellChat computed the interaction score for a signaling pathway by summing up the interaction strengths of all the L-R involved in that pathway. Based on these new signaling scores, dysregulated signaling were identified between control and KO groups using *rankNet* function (Wilcoxon test, p < 0.05), with a tolerance > 0.05 for the differences in the relative contributions.

## Results

### Clustering and annotation of cell types

To model CHD8 functions in early neural differentiation, we generated human cerebral organoids from two control iPSC lines (“iPS1” and “iPS2”) and two derived lines (“KO1” and “KO2”) with a copy of the *CHD8* gene edited by CRISPR-Cas9 (*CHD8*^+/-^ referred as heterozygous *CHD8* knockout (KO) for simplicity). The generation and characterization of the lines were described in our previous studies, which also confirmed a reduction of CHD8 protein in the KO lines ^13,14^. The transcriptomic changes of *CHD8* KO were also analyzed in neuron progenitors, differentiated neurons, and cerebral organoids by bulk RNA-seq analysis ^13,14^. In this study, cerebral organoids after 50 days of neural induction were harvested for scRNA-seq analysis.

After low quality cells were removed, we obtained an average of 14,285 cells per sample, with an average of 25,398 reads per cell and 2,184 genes per cell (**Supplementary Table S1**). The cell level statistics was similar in all four samples. The data were then analyzed and integrated with RISC software, which was specifically designed for batch correction and integrating cells with expected treatment effects ^33^. Additional filtering in RISC, to retain high quality cells, yielded an average number of 12,287 cells per sample, with maximal (13,723) and minimal (10,123) cells in KO2 and iPS2, respectively. Louvain clustering of the integrated data yielded 21 cell clusters, as visualized by dimension reduction and uniform manifold approximation and projection (UMAP) (**Figure 1A**). Differential analysis between one cluster and the remaining cells identified marker genes expressed highly in each of the 21 cluster (**Figure 1B,C**; **Supplementary Table S2**), which were used for cell type annotation, together with a set of markers reported in previous scRNA-seq studies of brain organoids (**Figure 1D; Supplementary Figure S1**) ^28,38,46^.

**Figure 1:**
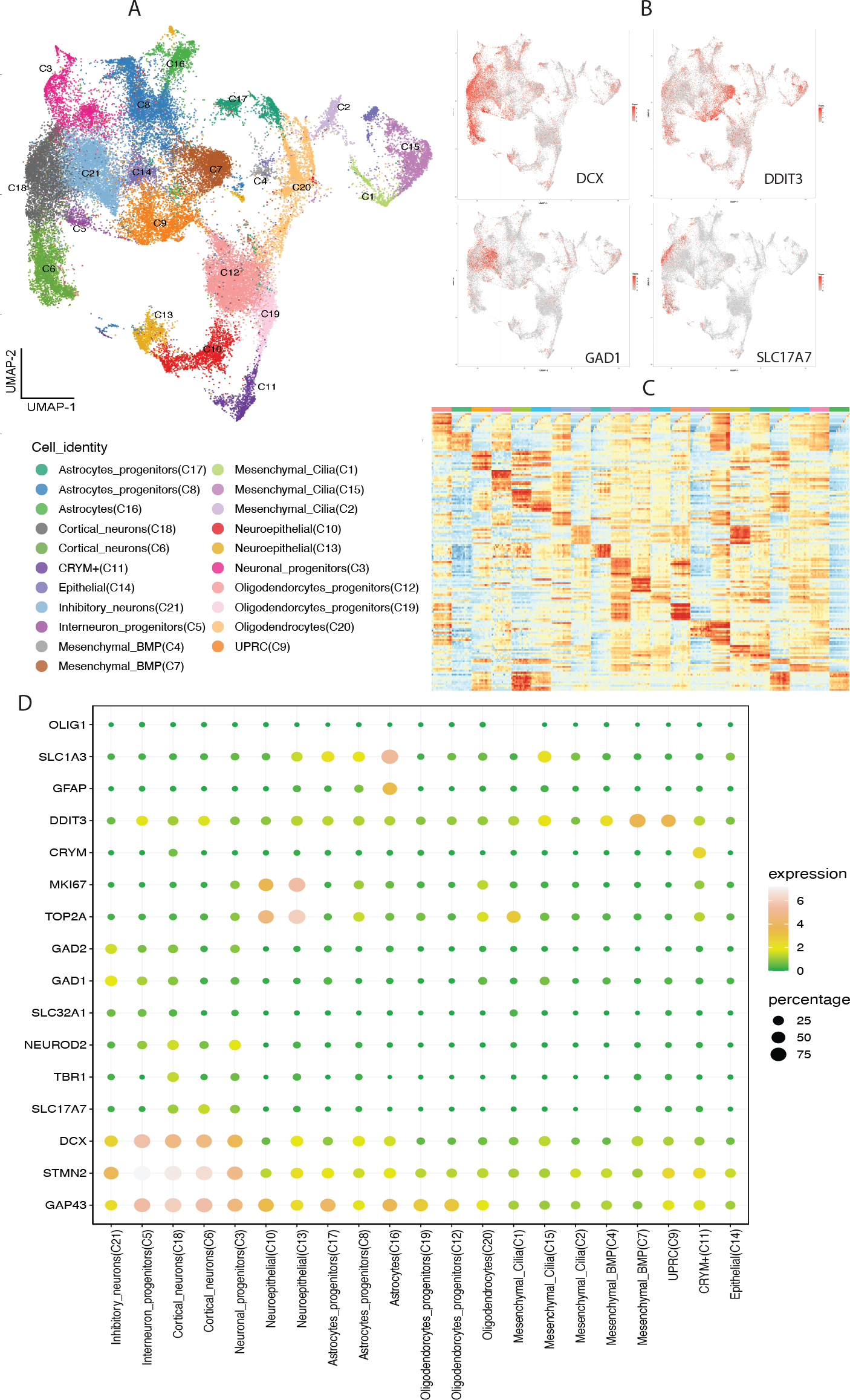
Cell clustering and annotation of the integrated scRNA-seq data. A: Uniform manifold approximation and projection (UMAP) of integrated data revealing 21 distinct cell clusters. B: UMAP plots for the expression of selected markers. C: Heatmap for the expression of top 10 markers computed for the 21 clusters (left to right, cluster 1 to 21). The names of the markers are not shown but can be found in **Supplemental Table S2**. D: Bubble plot for the expression of top gene markers identified and used for cell cluster annotation.

Five clusters showed high expression of neuronal *GAP43, STMN2, DCX* markers, while the remaining 16 clusters expressed high level of non-neuronal markers (**Figure 1D**). Three of the five neuronal clusters were annotated as excitatory cortical neurons (Cortical_neurons (C6), Cortical_neurons (C18)) based on their expression of *SLC17A7, TBR1* and *NEUROD2* markers, with one of them (Neuronal_progenitors(C3)) determined to be neural progenitor due to lower *SLC17A7* expression and their location in the early differentiation trajectories (see below; **Figure 2**). Two clusters of interneurons were found, based on *SLC32A1* expression (Interneuron_progenitors (C5)), with one of them (Inhibitory_neurons (C21)) expressing higher *GAD1* and *GAD2*, canonical inhibitory neuron markers.

**Figure 2:**
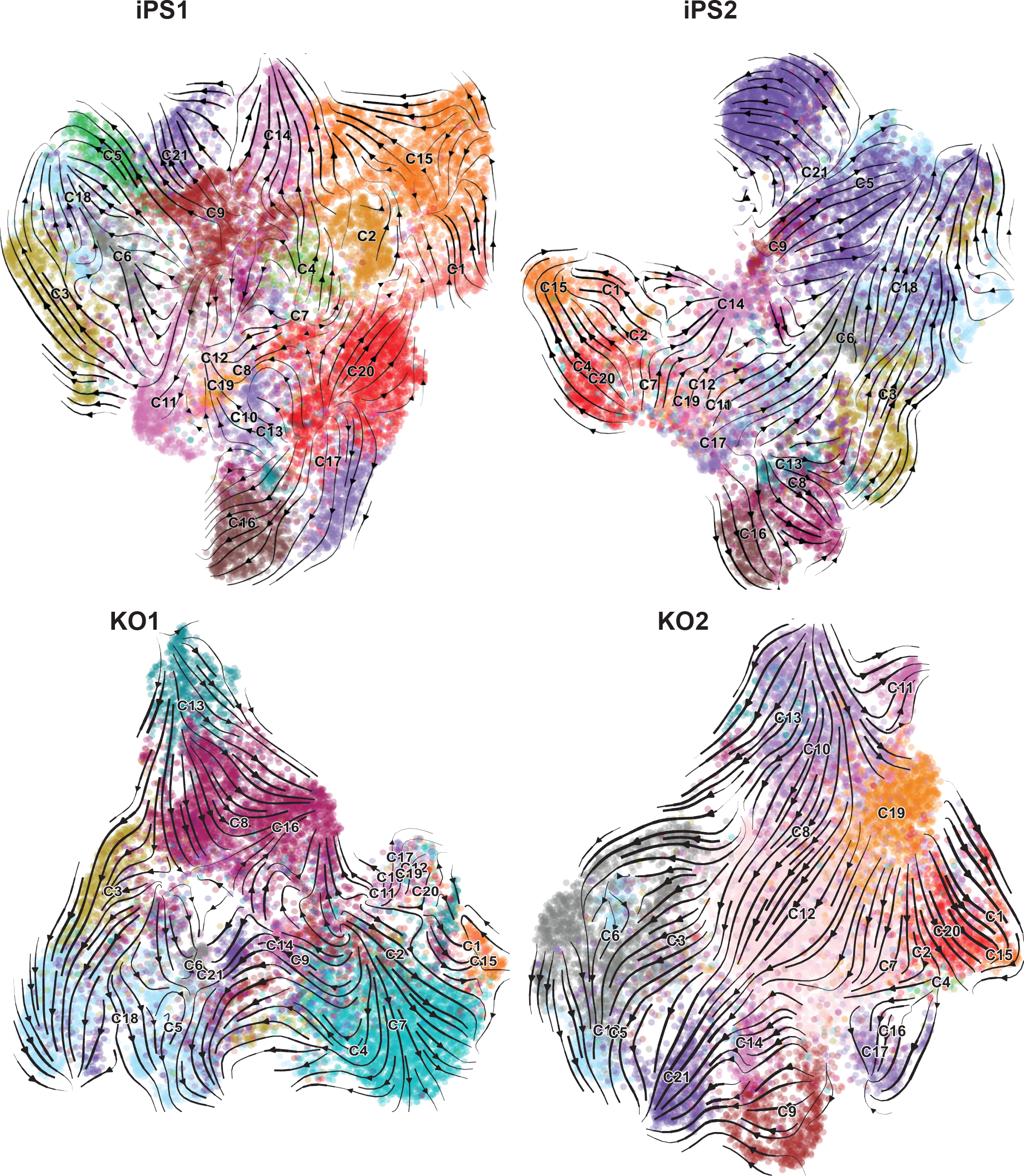
UMAP projection of cell differentiation trajectories predicted for each sample. RNA velocity trajectories were analyzed separately for each sample to avoid batch difference but cell clusters were taken the RISC integrated data.

For non-neuronal clusters, *TOP2A* and *MKI67* expression marked two neuroepithelial clusters (C10 and C13), while positive *CRYM* and *DDIT3* expression were used for identifying CRYM+ (C11) and UPRC (C9) cells, respectively ^38^. C9 markers were enriched for genes in the GO:0006986 (“response to unfolded protein”) supporting the cell identity. Three cluster of Astrocytes (C16), Astrocytes_progenitors (C8) and Astrocytes_progenitors (C17)) with *GFAP* and *SLC1A3* expression were identified, with the latter two expressing lower levels of these two markers than C16. RNA velocity, pseudotime trajectory analysis, and clustering based on scvelo dynamic data, indicated that C16 were more differentiated than C8 and C17 (**Figure 2**), thus the latter two were designed as astrocyte progenitors. Similarly, three clusters of oligodendrocytes: Oligodendrocytes (C20), Oligodendroytes_progenitors (C12), Oligodendrocytes_progenitors (C19)) were identified, based on positive, but at different levels, of *OLIG1* and *PDGFRA* expression, with the annotation supported by trajectory data. Mesenchymal cells were separated into five clusters, using markers described in a previous study ^38^. Marker genes for two of them showing significant enrichment in BMP pathway (Mesenchymal_BMP (C4), Mesenchymal_BMP (C7)) and the other three expressed high levels of cilium genes (Mesenchymal_Cilia (C15), Mesenchymal_Cilia (C2), Mesenchymal_Cilia (C1)). Lastly, one cluster, Epithelial (C14), showed enrichment of epithelial markers (**Figure 1**).

In short, scRNA-seq analysis indicate that the cerebral organoids from our differentiation protocols consist of a diverse array of neural progenitors, differentiating neurons, and non-neuronal cells (**Supplementary Figure S2**). Given that the organoids were harvested after 50 days, most of the differentiating cell types are likely in immature states, just starting to express cell type specific genes and thus making it challenging to make definitive cell type assignment. As such, our data and results should be considered as models suitable for addressing CHD8 roles in very early stages of neural differentiation, as described previously by Lancaster et al. ^31^.

### Neural population composition affected by CHD8 knockout

With the cell type identification, we set out to infer the differentiating trajectory relationship of cells in each sample. We performed trajectory analysis based on RNA velocity and gene expression similarity between cell clusters, using CellRank ^37^. RNA velocity predicts cell differentiation direction based on the ratio of splicing to unspliced RNAs ^35^. Surprisingly, our scRNA-seq reads captured a high number of intronic reads, up to 45% of unspliced reads in the iPS2 sample (**Supplementary Figure S3**), making it easy to perform RNA velocity analysis. In terms of cell types, reads for Epithelial (C14) and Inhibitory_neurons (C21) had the highest percentages of unspliced reads. On the other hand, Mesenchymal_Cilia (C2) showed the lowest proportions of unspliced reads.

We projected the inferred cell differentiation trajectory results to UMAP plots (**Figure 2**). The trajectories look largely similar for the four samples, suggesting that CHD8 reduction did not fully block neural differentiation potential, consistent with previous studies ^28,29^. In three of the four samples, Cortical_neurons (C18), Inhibitory_neurons (C21) and Astrocytes(C16) were identified as terminal states, while Epithelial (C14) cells as alternative terminal states for iPS1 and iPS2. Consistent with the expression analysis of marker genes (**Figure 1**), progenitors of Cortical neurons, Astrocytes and Oligodendrocytes were located to the intermediate states of differentiation in the UMAPs. The analysis also identified Neuroepithelial (C13) cells and UPRC (C9) cells as initial or early states in all samples, while Oligodendrocytes_progenitors (C19) cells were considered as an additional initial state for KO2 sample. Overall, the trajectory analysis helps to correctly align cell types to the known neural differentiation process.

We next studied if the cell population differences between *CHD8* KO and control samples could identify the points where differentiation is most regulated by CHD8, i.e., altered by *CHD8* KO. We used miloR, which applies KNN graph based differential abundance analysis on data with replicates and is more robust than many other methods, such as simple proportion test of cell numbers in individual clusters ^34^. We found that most of the cell types showed a decrease of abundance in the *CHD8* KO samples (**Figure 3**). The decrease was highly significant for the Oligodendrocytes (C20), Neuronal_progenitors (C3), Mesenchymal_Cilia (C15), Astrocytes (C16) and Astrocytes_progenitors (C17). On the other hand, Mesenchymal_BMP (C7) cell showed the highest increase in the KO samples. When the trends in all the detected cell types, especially the neural cells, were considered together, the data suggest that neuroepithelia (early progenitors C10 and C13) and cortical neurons (C6 and C18) were proportionally more abundant in the *CHD8* KO samples than controls, while inhibitory neurons were less abundant, suggesting that CHD8 plays important roles in neuronal fate specification. If confirmed in future study, the data points to a scenario that early neurogenesis (e.g., lineage specification) may decelerate in *CHD8* KO but the intermediate progenitors (e.g., C13), once produced, were accelerated to differentiate. This analysis also suggests that CHD8 could have important roles in glial cell differentiation, because astrocytes and mesenchymal cells were reduced in *CHD8* KO.

**Figure 3:**
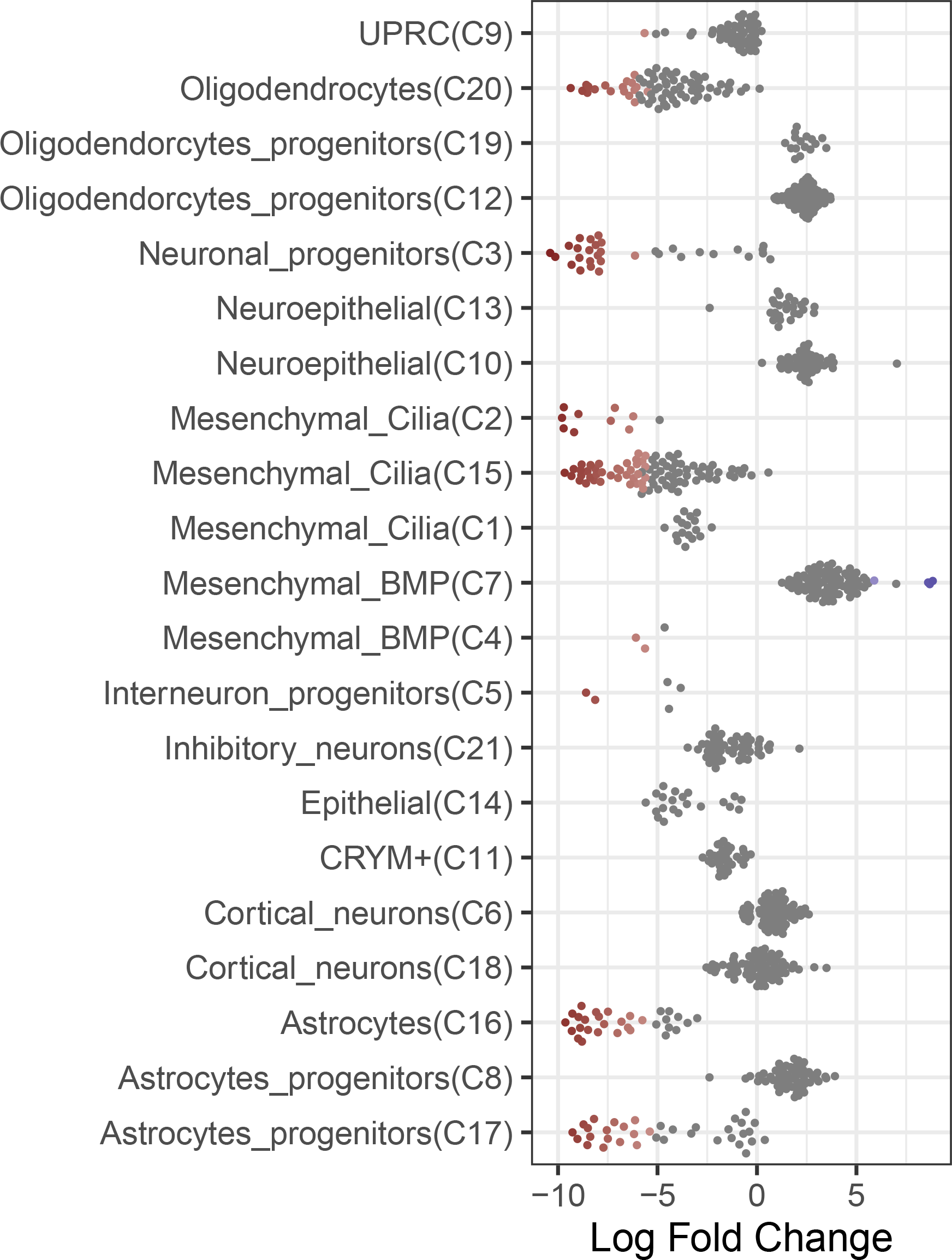
Beeswarm plot showing distribution of log fold change in the abundance of cell types. Individual dots represent neighborhoods containing cells from different cell type clusters, with blue for significantly higher abundance in KO and red for significantly higher abundance in control samples.

Taken together, this analysis suggests that CHD8 plays important roles in both neural and non-neural cell fate specification, but the perturbation from its reduction may be subtle. The effects seem to be more robust in previous reports that used organoids at later stages of differentiation^28,29^, when the perturbed phenotypes become more obvious.

### Effects of CHD8 haploinsufficiency on cell type specific gene expression

We next performed differential expression analysis between *CHD8* KO and control samples, for each of the cell cluster independently. We found a total of 4,913 unique genes affected by CHD8 haploinsufficiency in at least one of the cell types (**Supplementary Table S3**). Mesenchymal_BMP (C7) had the most differentially expressed genes (DEGs) (n=1,005), while Cortical_neurons (C6) had the least (n = 35), indicating that CHD8 may have more important roles in some cell types than others. We carried out Gene Ontology (GO) enrichment analysis of the DEGs, and the top five most enriched terms for each of the cell types were shown in **Figure 4**. The results demonstrated that GO terms such as synapse assembly, nuclear division, forebrain development, neuron projection guidance or axon guidance, were enriched in the DEGs detected in multiple neuronal cell types. For non-neuronal cell types, the enriched terms were related to axoneme assembly, central nervous system, neuron differentiation, cilium organization, extracellular matrix organization, modulation of chemical synaptic transmission or synapse assembly. The ECM difference was also observed in our previous bulk RNA-seq studies, while the axoneme and cilia formation abnormalities have been linked to the functions of ASD risk genes ^47–51^ (https://doi.org/10.53053/CMZZ2213).

**Figure 4:**
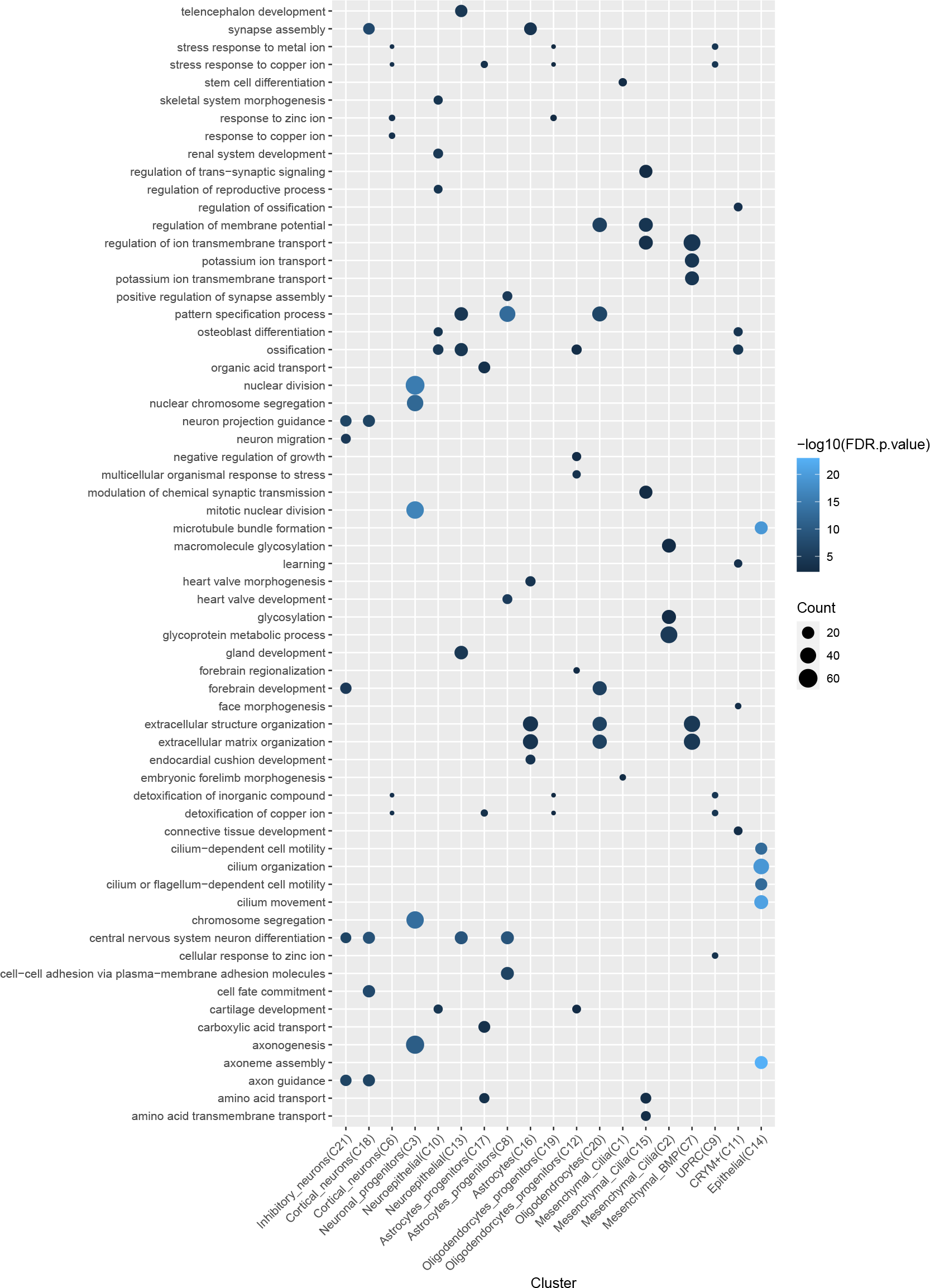
Dot plot showing the enriched GO terms for the dysregulated genes in *CHD8* KO.

In addition to GO term enrichment analysis based on DEGs selected with thresholds, we applied Gene Set Enrichment Analysis (GSEA) to genes expressed in each cell type after they were ranked by expression changes between *CHD8* KO and controls. From this analysis, inhibitory neurons showed upregulation of genes enriched in GO terms related to alcohol metabolomics, cell junction, lipid biosynthesis, regulation of synapse assembly or steroid biosynthesis (**Figure 5A)**. In contrast, most of the identified gene sets in cortical neurons showed a negative enrichment for sensory system development, nephron development or kidney and eye morphogenesis. Neuroepithelial cells showed downregulation for cell-cell adhesion, ensheathment of neurons, cerebral cortex development, glial cell differentiation and regulation of DNA binding and axonogenesis.

**Figure 5:**
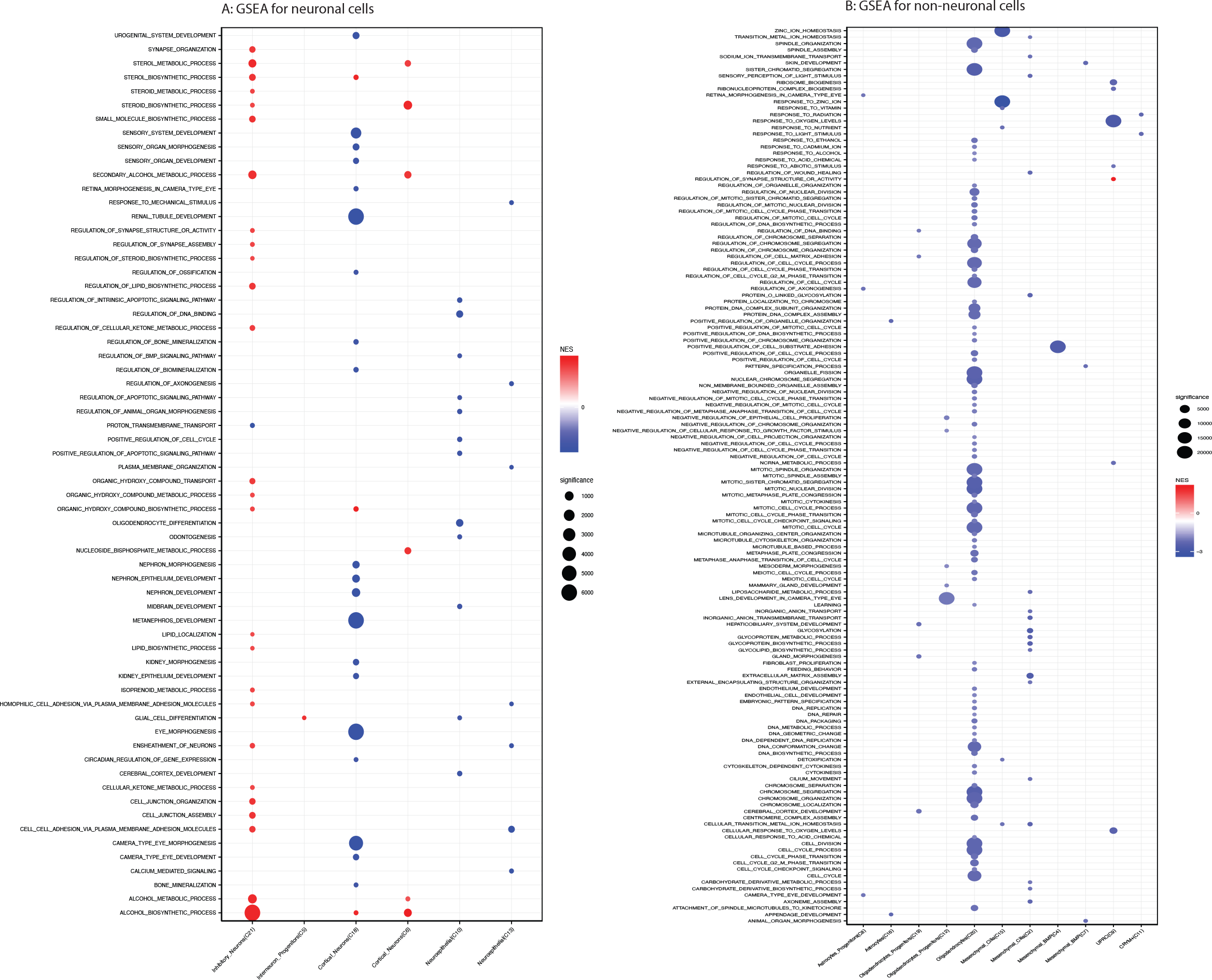
Dot plot showing enriched pathways from GSEA for neuronal (A) and non-neuronal (B) cell clusters, with red and blue for higher and lower activities in the *CHD8* KO samples, respectively.

Most of the enriched terms from GSEA for non-neuronal cells were found in the downregulated genes in *CHD8* haploinsufficiency, with Oligodendrocytes having the most identified enriched pathways (**Figure 5B**). Reduced expression genes in astrocytes showed an enrichment of eye/appendage development and regulation of axonogenesis. In mesenchymal cells, down-regulated genes were enriched in terms of axonome assembly, cilium movement, matrix assembly or cell substrate adhesion. CRYM+ and UPRC cells showed enrichment in response to radiation and light stimulus and response to abiotic stimulus and oxygens levels, respectively. Dysregulated genes in the oligodendrocytes were enrichment in cell cycle, chromosome organization, DNA repair, DNA replication, nuclear chromosome segregation or protein DNA complex assembly among others.

### Cell-type effects of CHD8 haploinsufficiency on autism risk

To explore how the dysregulated genes are related to autism and other neural disorder risk, we compared the cell type specific DEGs to gene sets associated with brain disorders (**Figure 6**), including genes associated with ASD, schizophrenia (SZ), bipolar disorder, intellectual disability (ID), and attention deficit hyperactivity disorder (ADHD) ^41^. Most of the cell types showed an enrichment of genes in the ASD_SFARI, ASD_AUTISMKB, ADHD and BIPOLAR databases, while only Cortical_neurons (C18 and C6) showed an enrichment in genes associated with Schizophrenia (SZ_GWAS). Interestingly, dysregulated genes in the Mesenchymal_BMP (C4) cluster were overrepresented with genes associated with intellectual disability (ID_CNV). It is interesting to see a strong enrichment with ASD genes but nearly no enrichment for SZ and ID risk genes. In total, 157 of the ASD SFARI genes showed differential expression in at least one of the 21 clusters, with the most in the Neuron progenitor cluster (C3; n=43) and Astrocyte (C16; n=34), including *NRXN2/3, SHANK2 and SMARCA2*. The altered expression of selected genes is shown in **Supplementary Figure S4**.

**Figure 6:**
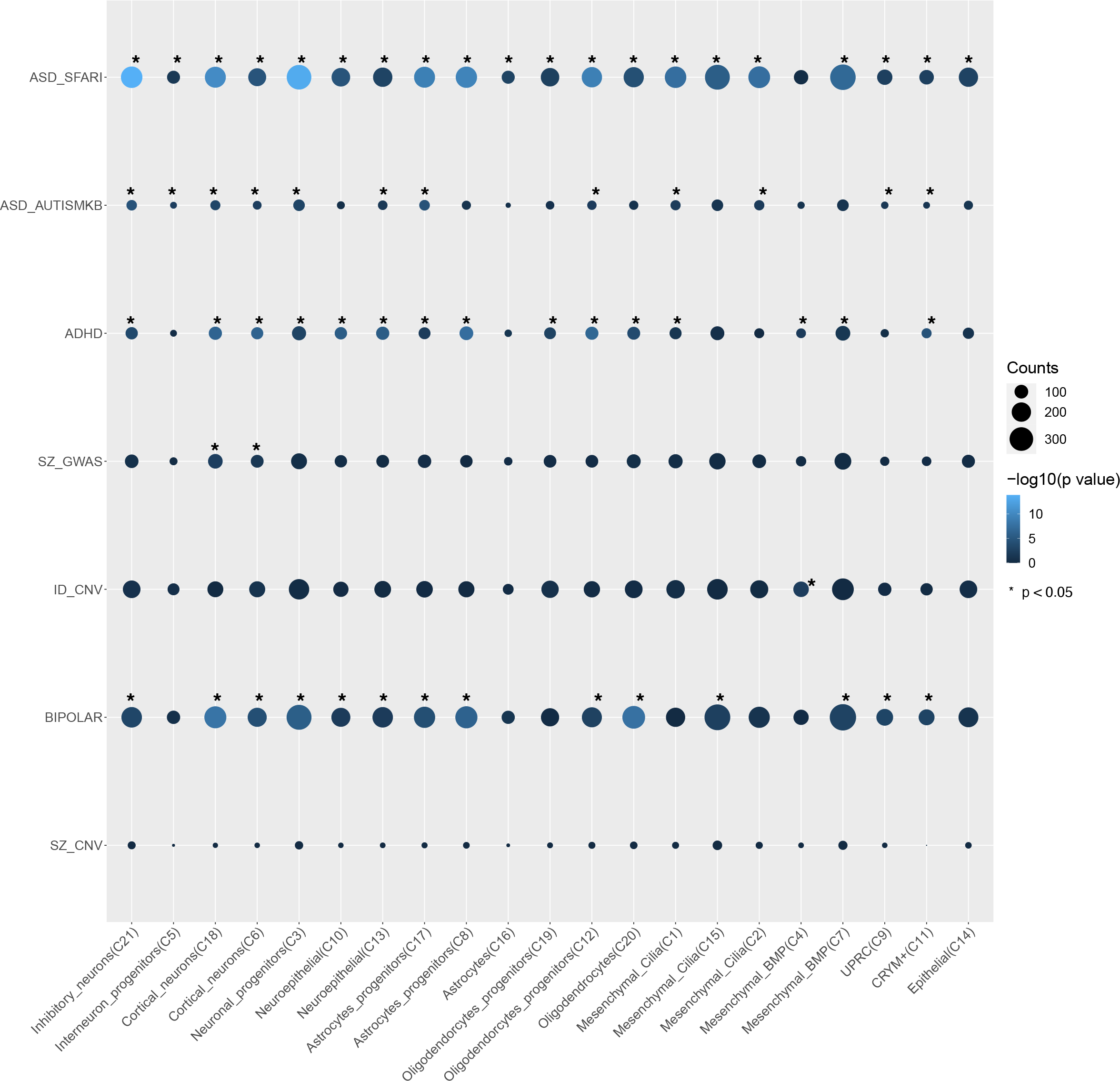
Dot plot showing the enrichment between dysregulated genes and lists of genes implicated in several brain disorders. The p-values were from over-representation analysis by hypergeometric test.

### Effects of CHD8 haploinsufficiency on cell-cell communication

In addition to studying cell population changes and cell type specific effects, scRNA-seq data are also valuable for investigating cell-cell communications across different cell types ^41^. We thus applied CellChat ^45^ and identified 12,763 ligand-receptor (L-R) interactions in the two iPS control samples and a slightly fewer interactions in in the *CHD8* KO samples (11,490) (**Figure 7**). Astrocytes (C16) showed the highest decrease in the number of interactions, specially, their interactions with Cortical neurons and Oligodendrocytes (**Figure 7A, Supplementary Figure S5**). On the other hand, Mesenchymal cells showed increased numbers of interactions. The interaction strengths between most pairs of cell types were also reduced, with an overall strength of 608.693 in the KO compared to 867.701 for the iPS samples. As shown in **Figure 7B** and **Supplementary Figure S6**, the majority of the cell types exhibited a decreased interaction strength, with only Mesenchymal cells having an increase. The altered L-R interactions converged to 55 signaling pathways (**Figure 7C**), with 3 (VEGF, THBS, NEGR) detected only in the control iPS samples and 4 (EDA, AMH, CHEMERIN, TAC) seen only in the KO samples, some of these altered signaling may be important for non-neural cells’ supports to the growth of neural cells. Overall, 30 signaling pathways showed a decreased activity in KO samples, while 25 displayed the opposite trend. 41 of the 55 were determined by the rankNet function as statistically significantly different between control and KO samples (**Figure 7C**). Interestingly, changes in the CNTN and NRXN signaling network (**Figure 7D,E**) were also observed in the brains from ASD subjects types ^41^. Details of the L-R interactions and cell-cell signaling differences are described in **Supplementary Table S4**. The changes involve some well-known ligands and receptors, including CNTN2, CNTNAP2, FGF, WNT, NRXN1, and RELN, which were also found in our differential expression analysis (**Supplementary Figure S7**). Additionally, a change in Wnt/μ-catenin signing (**Figure 7F)** upon *CHD8* reduction was also identified in our previous study ^13,14^.

**Figure 7:**
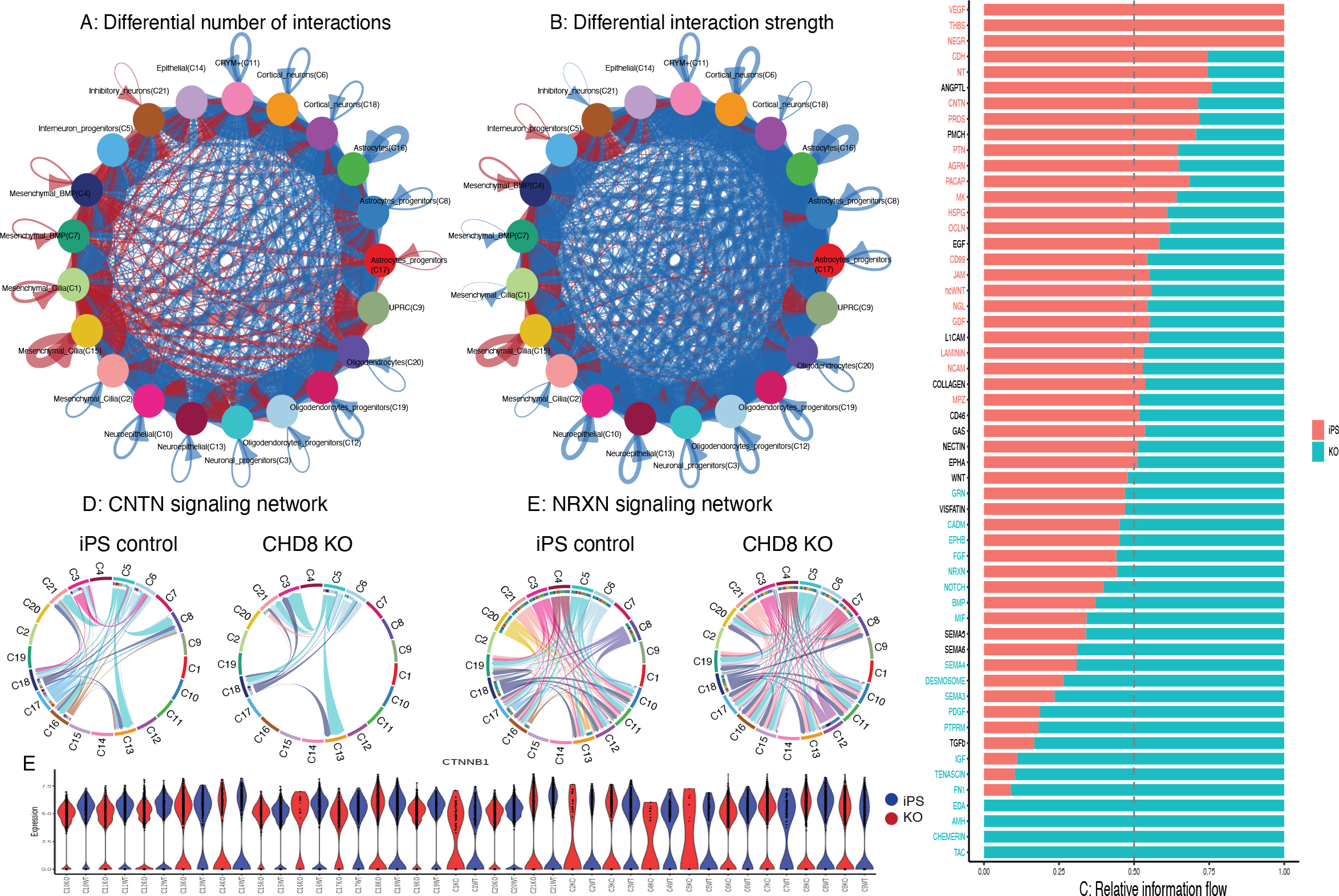
Change in cell-cell communications between *CHD8* KO and controls. A) Difference in the total numbers of L-R interactions in KO vs control samples. B) Difference in the total strengths of L-R interactions in KO vs control samples. C) Significantly altered cell-cell signaling pathways ranked by their differences in the overall information flow within the inferred networks between control and KO samples. D,E) Chord diagrams plotting connection difference for CNTN (D) and NRXN (E) network. F) Violin plot showing the expression of *CTNNB1*.

## Discussion

After ASD risk genes are identified, deciphering their functional roles is a big challenge. *CHD8* has been extensively studied since its discovery as a risk gene in 2012, with more than 200 publications. Here, we apply cerebral organoid and scRNA-seq technologies to investigate its function in early neural development and patterning at single cell resolution. Similar studies have been reported recently ^28–30^. In one, two other ASD risk genes, *SUV420H1* (also known as *KMT5B*) and *ARID1B* were studied together with *CHD8*, and all were found to play critical roles in neural development, as their haploinsufficiency conferred asynchronous development of GABAergic neurons and deep-layer excitatory neurons ^29^. The effects, however, were dependent on iPSC lines and neural differential stages, with two of the four *CHD8* lines showing an increase of GABAergic neurons and their progenitors in 3.5 and 6 months of post neural induction. This is consistent with our previous report of an increased expression of inhibitory neuronal markers (e.g., *DLX1* and *DLX6-AS1*) in cerebral organoids generated by a different protocol ^52^. Two other *CHD8* lines, however, showed no significant difference in GABAergic interneurons ^29^. Similarly, Villa et al also found an increase in inhibitory neurons but a delay of excitatory neurons at day 60 and 120, leading to an imbalance of the two major neuron types^28^. Interestingly, they found that *CHD8* haploinsufficiency caused an increased proliferation of the organoids, starting from day 10 and persistent to day 60, with an expansion of neural progenitors. While these two reports and our current study all show a disruption in neural developmental trajectories and their dynamics, the effects on specific cell types seem different, suggesting the effects of *CHD8* haploinsufficiency are likely modulated by genomic contexts, differentiation protocols and stages of development. In addition to the alterations in individual neuron cell types, our study also identified perturbation in non-neuronal cells and cell-cell communications in the cerebral organoids. Among the differential cell-cell interaction signaling, some are important for angiogenesis or vascular network patterning, for example, VEGF, THBS, and FN1. Alternation of such signaling may make the *CHD8* deficient cells more prone to environmental changes, leading to development disruptions. Along the same line, this may be a reason why variations of the KO effects were observed in different CHD8 studies, because the results may be affected by differentiation protocols and even media. Although more studies are needed to follow up these findings, it is conceivable that the increase of ASD risks in individuals with functional mutations can come from non-neuronal cells too. It will be interesting if such intracellular signaling involve the cilia on the cells that we annotated as Mesenchymal_Cilia.

Two main limitations of our current study are small sample sizes and a single time point. In addition, as the cerebral organoids were at a very early stage of differentiation, the separation of some cells into clusters may reflect more on the difference in transcriptomic states rather than cell types. Nevertheless, our scRNA-seq findings and others together provide a broad survey of the cell type dependent and context-specific roles of CHD8, indicating that the impact of *CHD8* mutations need more extensive analyses to map out the full functional spectrum.

## Competing interests

None to declare.

## Funding

This work was supported in part by grants to The Rose F. Kennedy Intellectual and Developmental Disabilities Research Center (RFK-IDDRC) from the Eunice Kennedy Shriver National Institute of Child Health & Human Development (NICHD) at the NIH (U54HD090260; P50HD105352).

## Authors’ Contributions

M.A. and Y.L. performed the bioinformatics analysis. E.P. and H.L. prepared the cerebral organoids and the scRNA-seq libraries. D.Z. and H.L. conceived of the experimental design. M.A., H.L. and D.Z. wrote the manuscript. D.Z. contributed to the data analytic plan and supervised the work. All authors read, edited, and approved the final manuscript.

## Acknowledgements

We would like to thank the supports from the Einstein Epigenomics Shared Facility, Einstein Computational Genomics Core, and Einstein High Performance Computing.

## Supplementary Tables

**Supplementary Table 1: Summary of the quality of the scRNA-seq data Supplementary Table 2: Marker genes identified for each cell cluster**.

**Supplementary Table 3: Lists of DEGs within each cluster between CHD8 KO and control samples**.

**Supplementary Table 4: Results from cell-cell interaction analysis using CellChat**. Table S4A lists L-R interactions in the iPS control dataset, Table S4B lists L-R interactions in the KO dataset, Table S4C lists signaling pathways in iPS control dataset, Table S4D lists signaling pathways in KO dataset, and Table S4E describes the relative contribution of signaling pathways within each condition.

## Supplementary Figures legends

**Supplementary Figure S1. Heatmaps for the expression pattern of selected marker genes used in previous publication**.

**Supplementary Figure S2: Dendrogram for cluster similarity based on mean gene expression**.

**Supplementary Figure S3: Proportion of spliced/unspliced reads for each cell cluster in the four samples**.

**Supplementary Figure S4: Violin plots showing expression of genes implicated in ASD**

**Supplementary Figure S5: Heatmap showing differences in cell-cell communication numbers between control and KO samples**.

**Supplementary Figure S6: Heatmap showing differences in cell-cell communication strengths between control and KO samples**.

## Notes

### Competing Interest Statement

The authors have declared no competing interest.

