## Supplemental Figures for "Molecular and network disruptions in neurodevelopment uncovered by single cell transcriptomics analysis of *CHD8* heterozygous cerebral organoids"

**Figure S1. Heatmaps for the expression pattern of selected marker genes used in previous publication.**

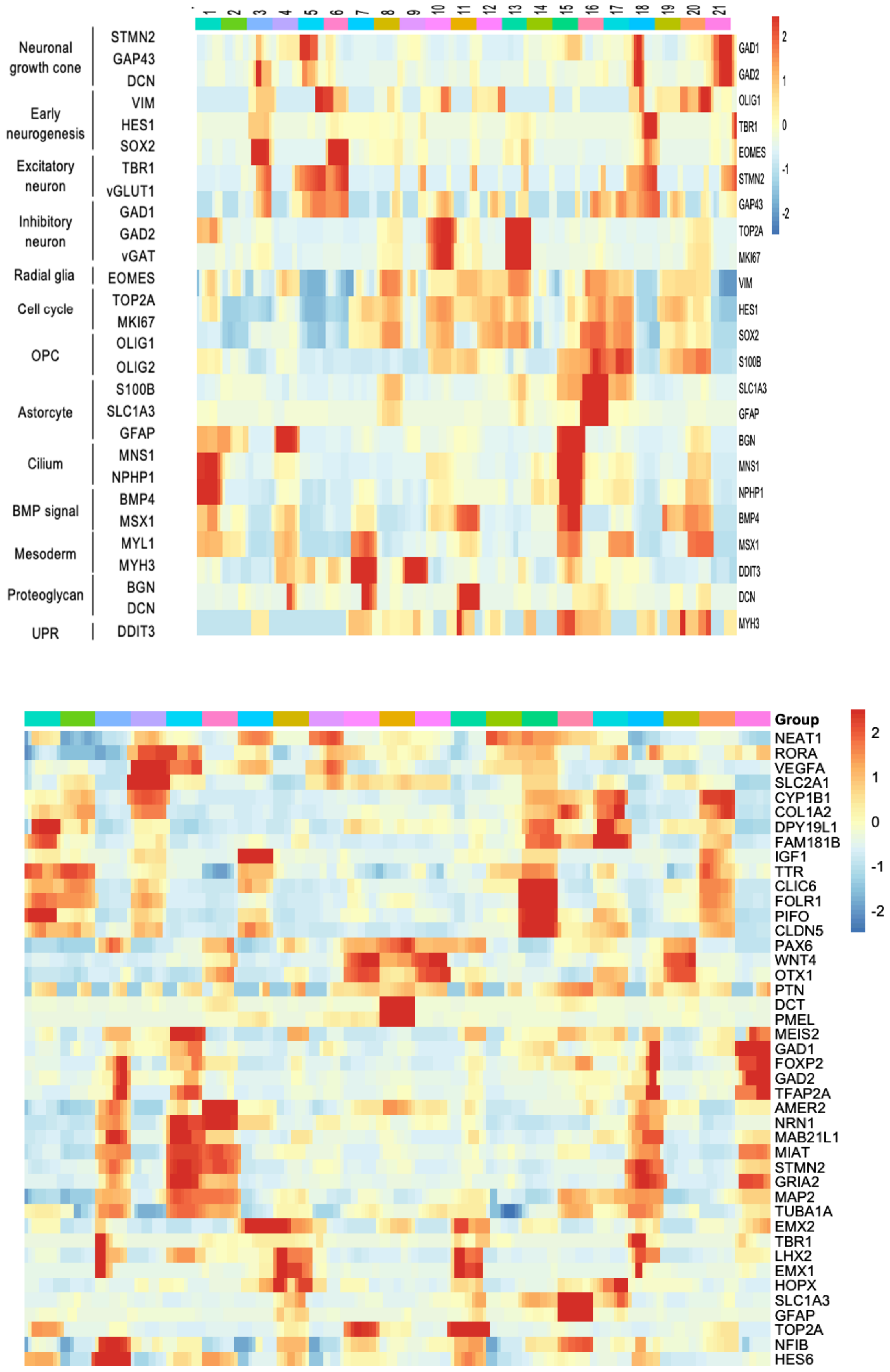

**Figure S2. Dendrogram for cluster similarity based on mean gene expression**

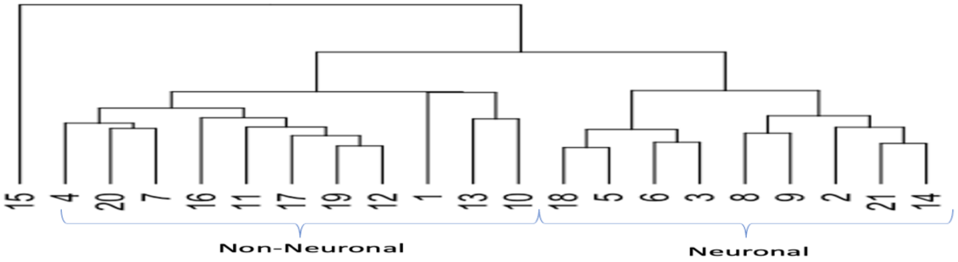


**Figure S3. Proportion of spliced/unspliced reads for each cell cluster.**

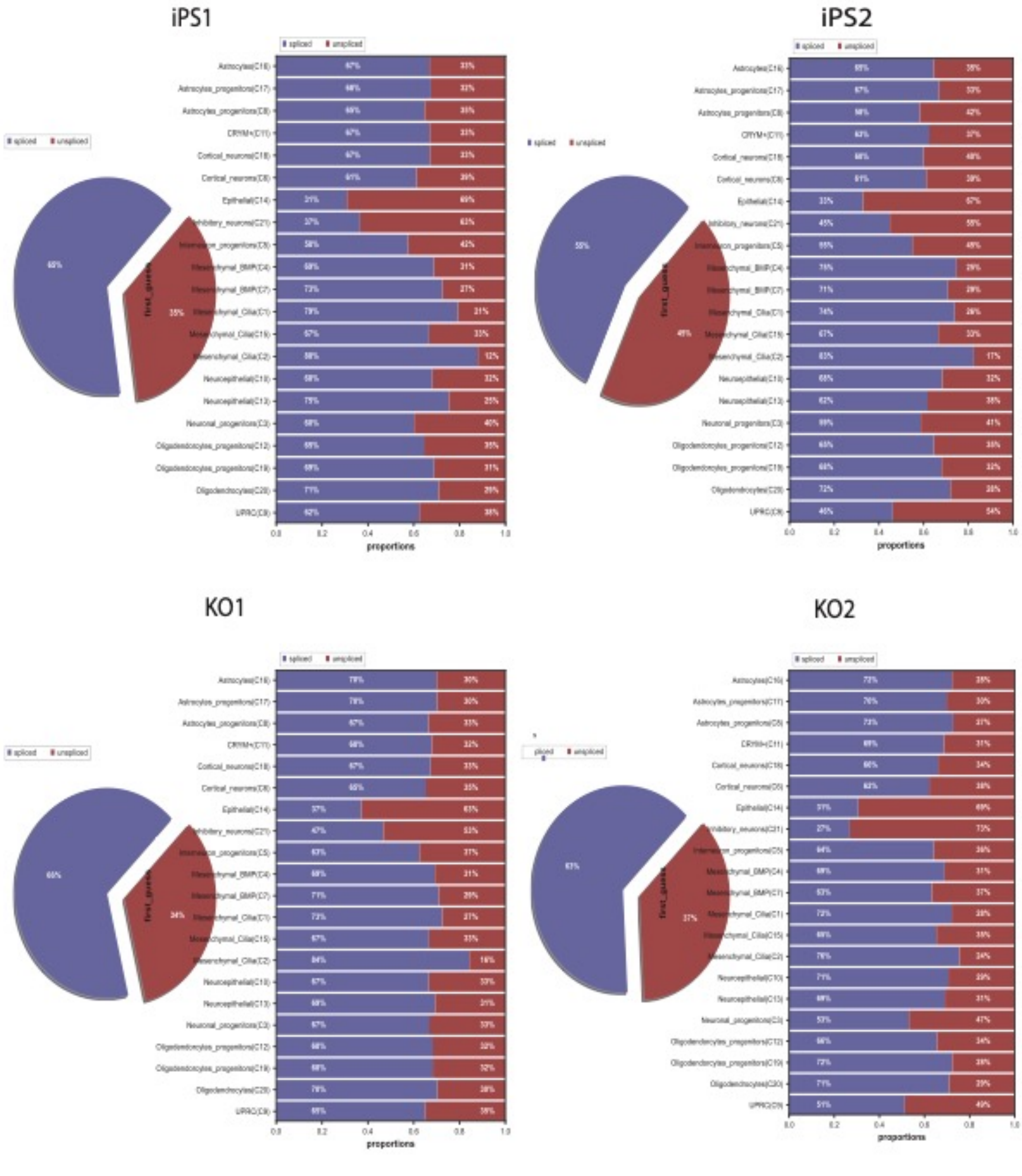

### Figure S4: Violin plots showing expression of genes implicated in ASD

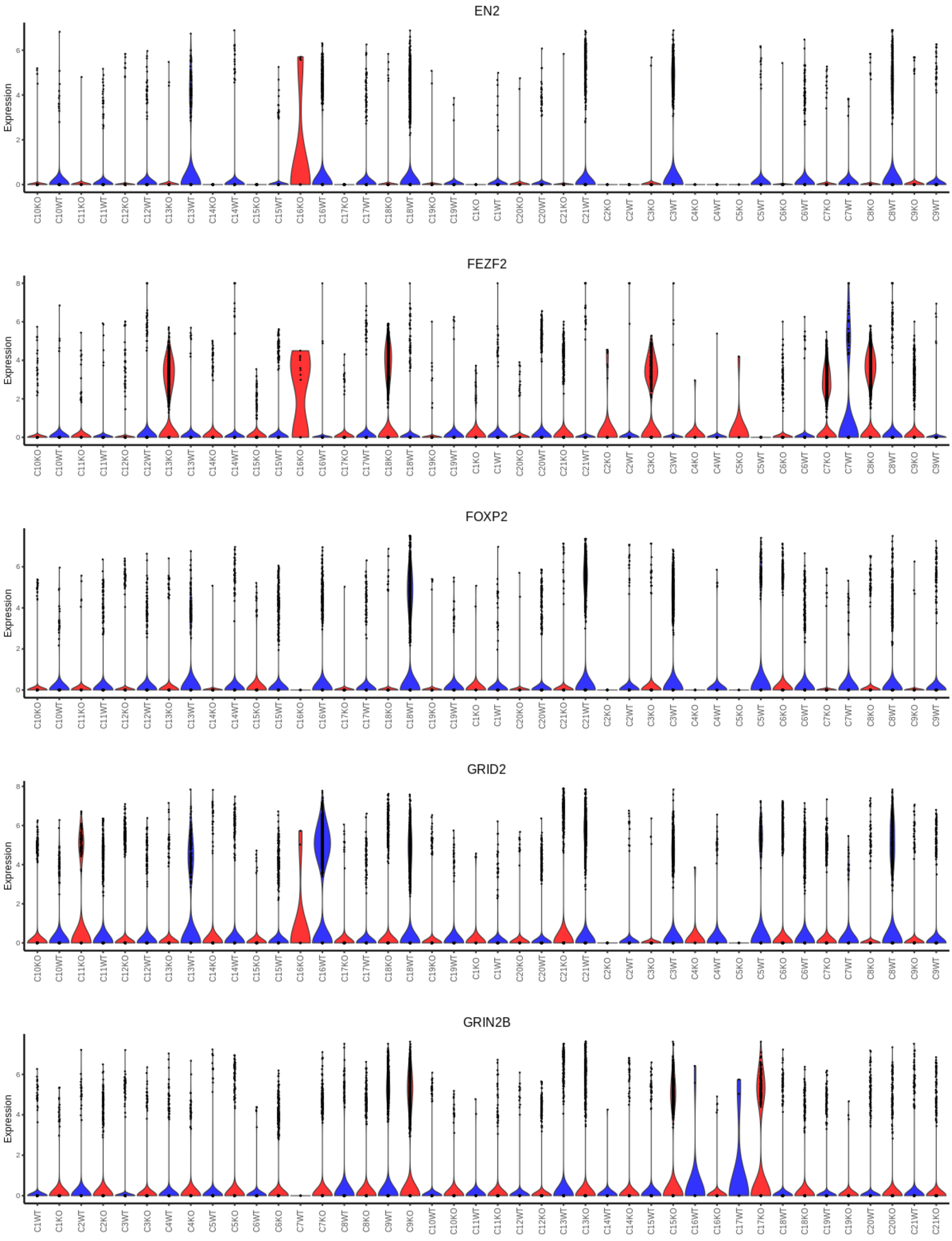

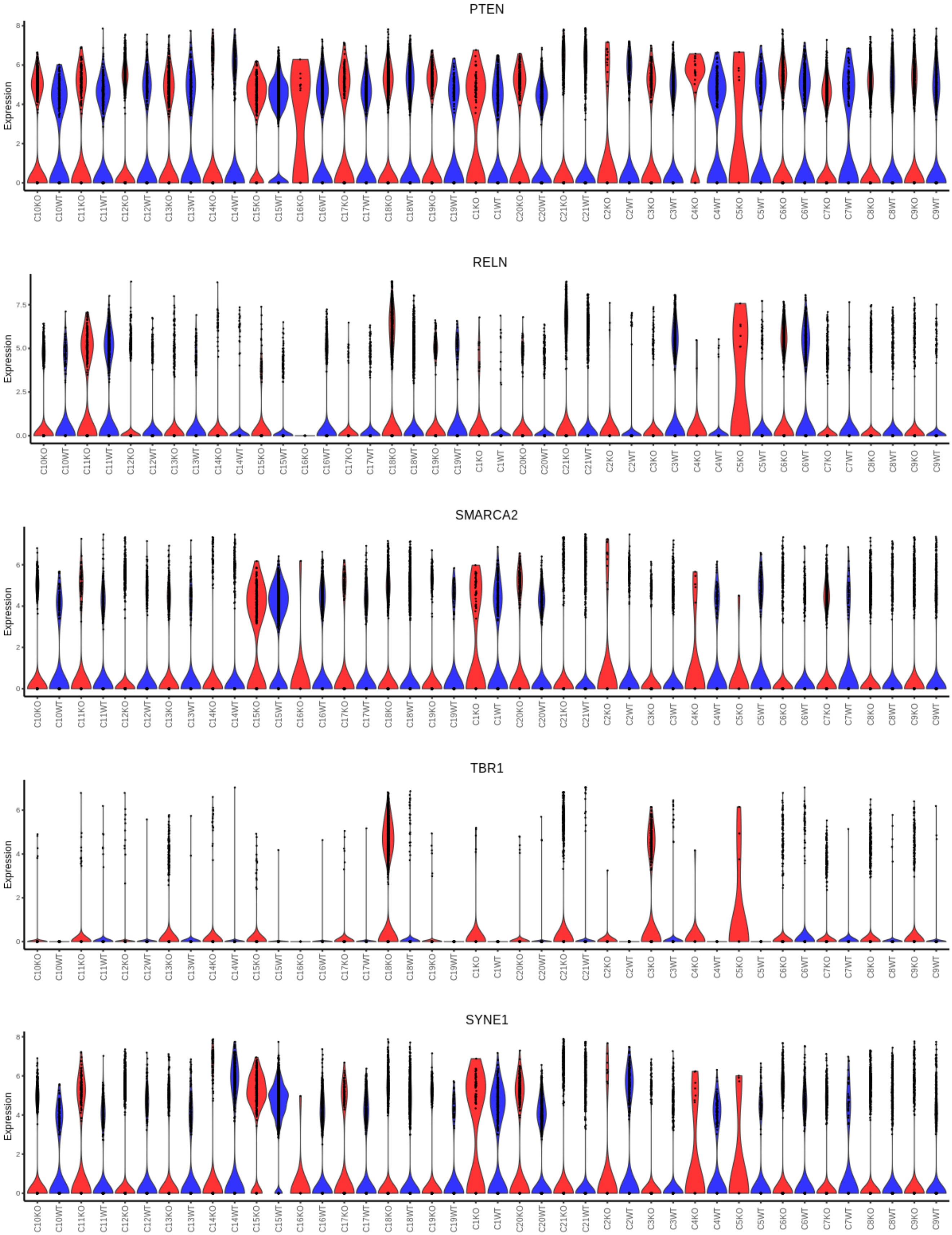
PTEN
Expression
0 2 4 6 8
C10KO C10WT C11KO C11WT C12KO C12WT C13KO C13WT C14KO C14WT C15KO C15WT C16KO C16WT C17KO C17WT C18KO C18WT C19KO C19WT C1KO C1WT C20KO C20WT C21KO C21WT C2KO C2WT C3KO C3WT C4KO C4WT C5KO C5WT C6KO C6WT C7KO C7WT C8KO C8WT C9KO C9WT
RELN
Expression
0.0 2.5 5.0 7.5
C10KO C10WT C11KO C11WT C12KO C12WT C13KO C13WT C14KO C14WT C15KO C15WT C16KO C16WT C17KO C17WT C18KO C18WT C19KO C19WT C1KO C1WT C20KO C20WT C21KO C21WT C2KO C2WT C3KO C3WT C4KO C4WT C5KO C5WT C6KO C6WT C7KO C7WT C8KO C8WT C9KO C9WT
SMARCA2
Expression
0 2 4 6
C10KO C10WT C11KO C11WT C12KO C12WT C13KO C13WT C14KO C14WT C15KO C15WT C16KO C16WT C17KO C17WT C18KO C18WT C19KO C19WT C1KO C1WT C20KO C20WT C21KO C21WT C2KO C2WT C3KO C3WT C4KO C4WT C5KO C5WT C6KO C6WT C7KO C7WT C8KO C8WT C9KO C9WT
TBR1
Expression
0 2 4 6
C10KO C10WT C11KO C11WT C12KO C12WT C13KO C13WT C14KO C14WT C15KO C15WT C16KO C16WT C17KO C17WT C18KO C18WT C19KO C19WT C1KO C1WT C20KO C20WT C21KO C21WT C2KO C2WT C3KO C3WT C4KO C4WT C5KO C5WT C6KO C6WT C7KO C7WT C8KO C8WT C9KO C9WT
SYNE1
Expression
0 2 4 6 8
C10KO C10WT C11KO C11WT C12KO C12WT C13KO C13WT C14KO C14WT C15KO C15WT C16KO C16WT C17KO C17WT C18KO C18WT C19KO C19WT C1KO C1WT C20KO C20WT C21KO C21WT C2KO C2WT C3KO C3WT C4KO C4WT C5KO C5WT C6KO C6WT C7KO C7WT C8KO C8WT C9KO C9WT

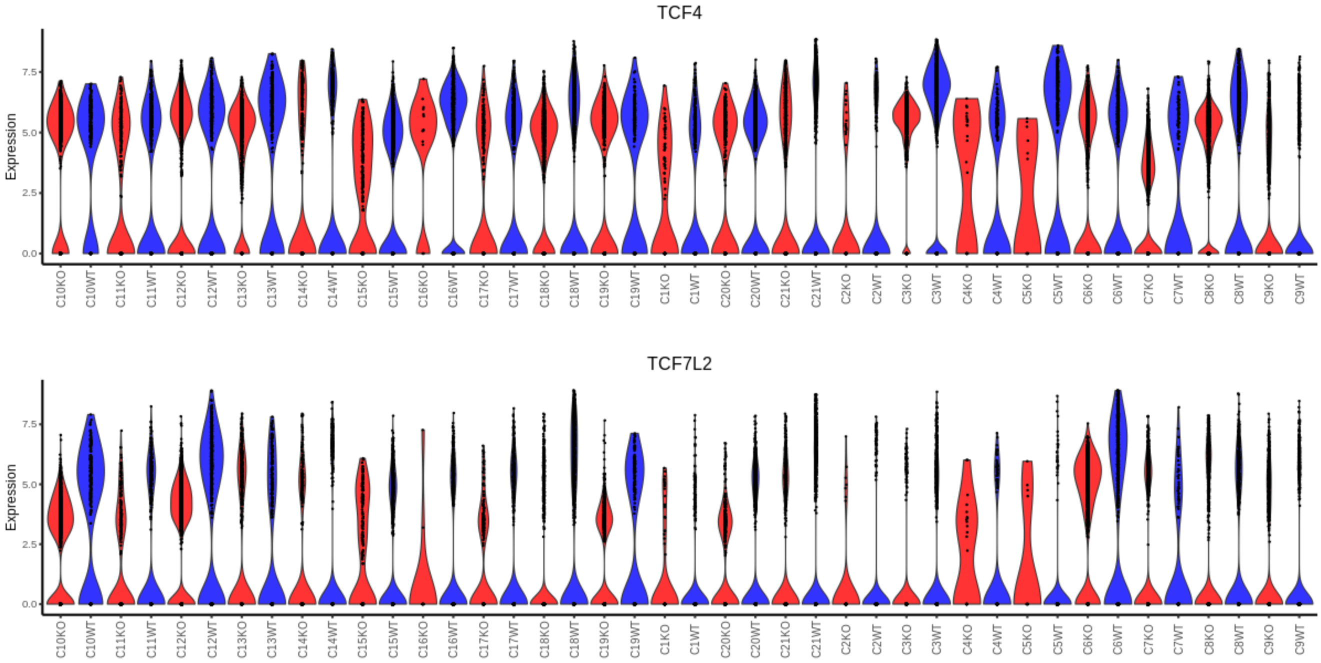


**Red: CHD8 KO cells, Blue: control cells**

**Figure S5. Heatmap showing cell-cell communication count differences between control and CHD8 KO samples.** The rows and columns indicate signaling sending and receiving cells, respectively.

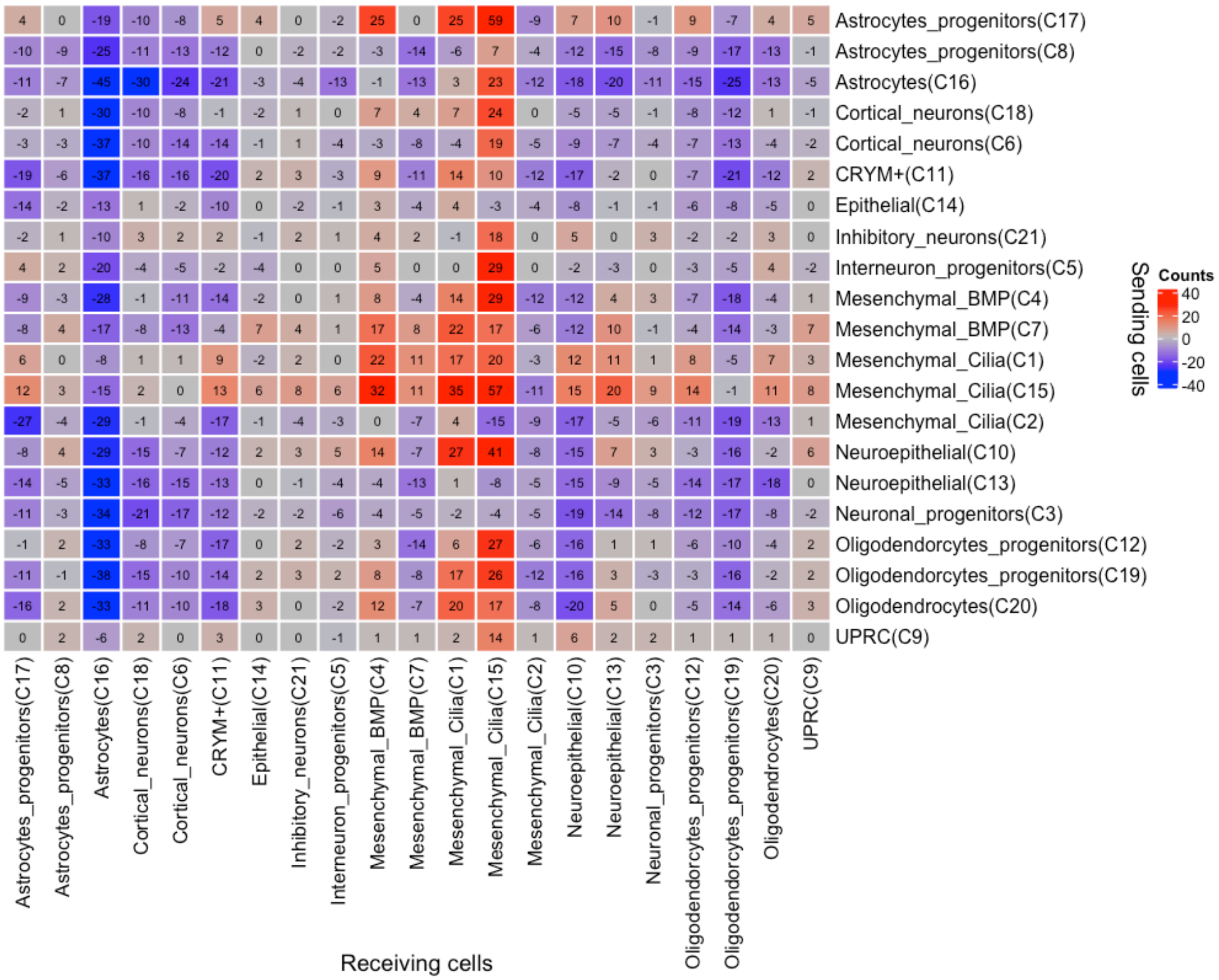

**Figure S6. Heatmap showing cell-cell communication strength differences between control and CHD8 KO samples.** The rows and columns indicate signaling sending and receiving cells, respectively.

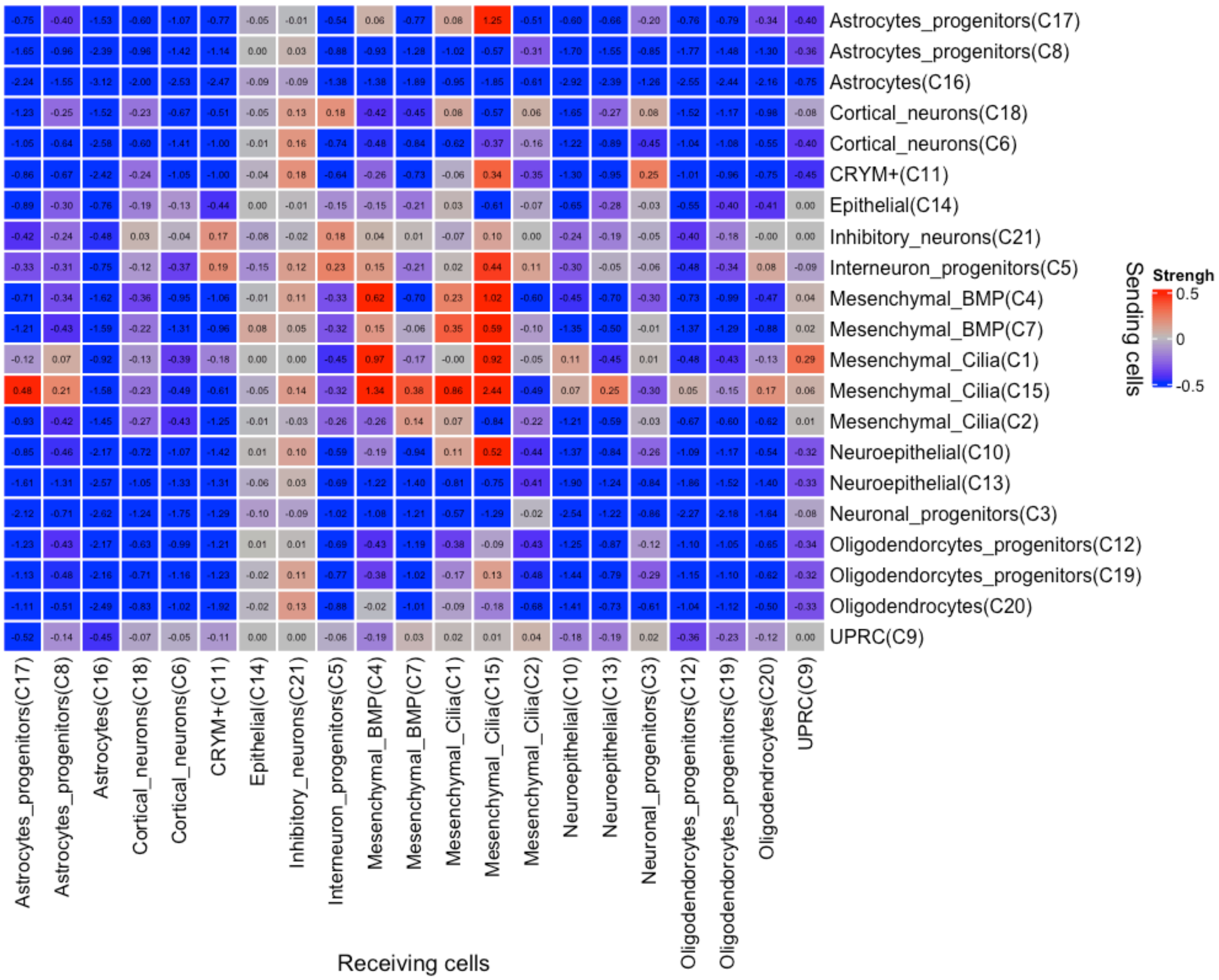

### Figure S7. Violin plots showing expression of genes encoding ligands or receptors

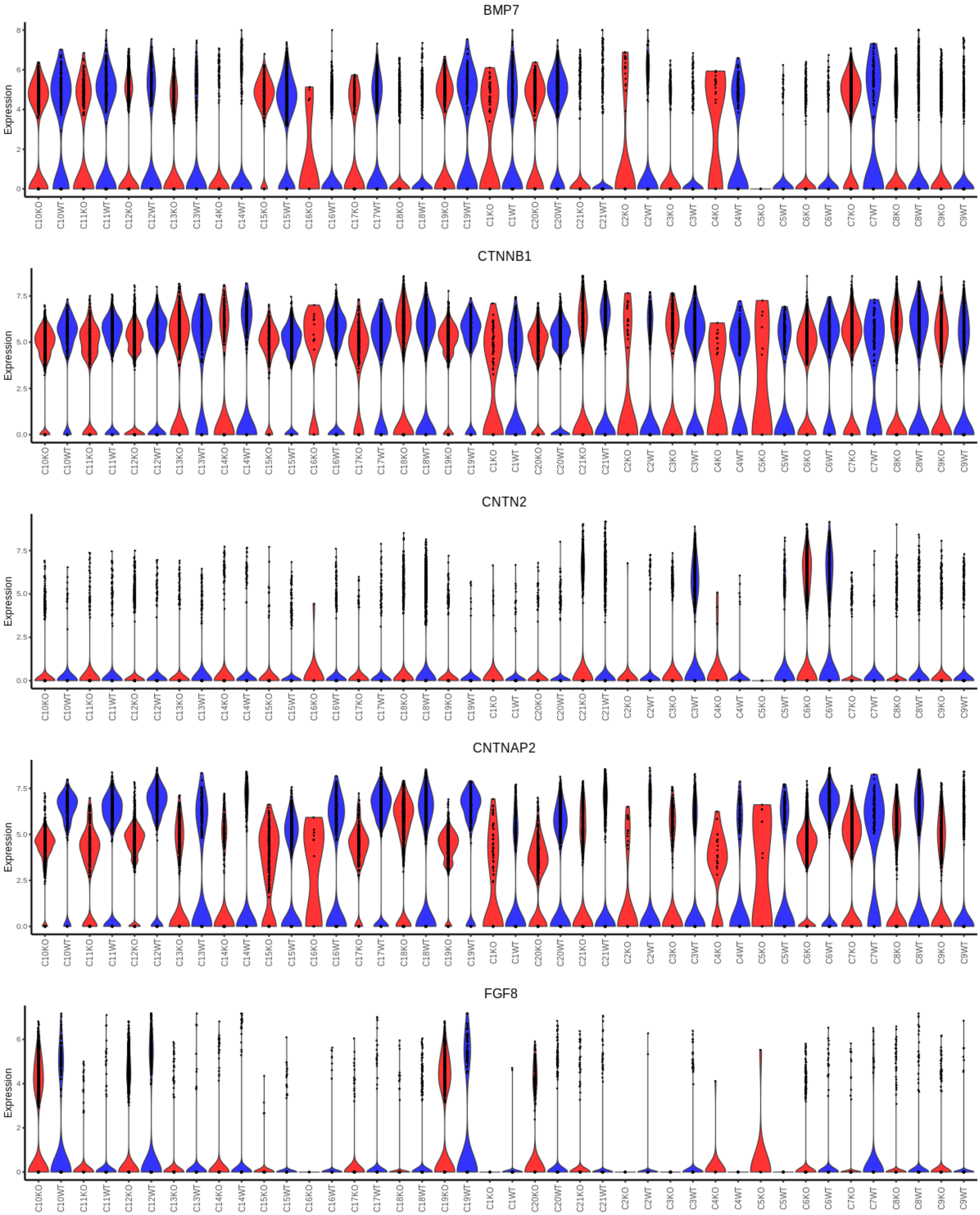

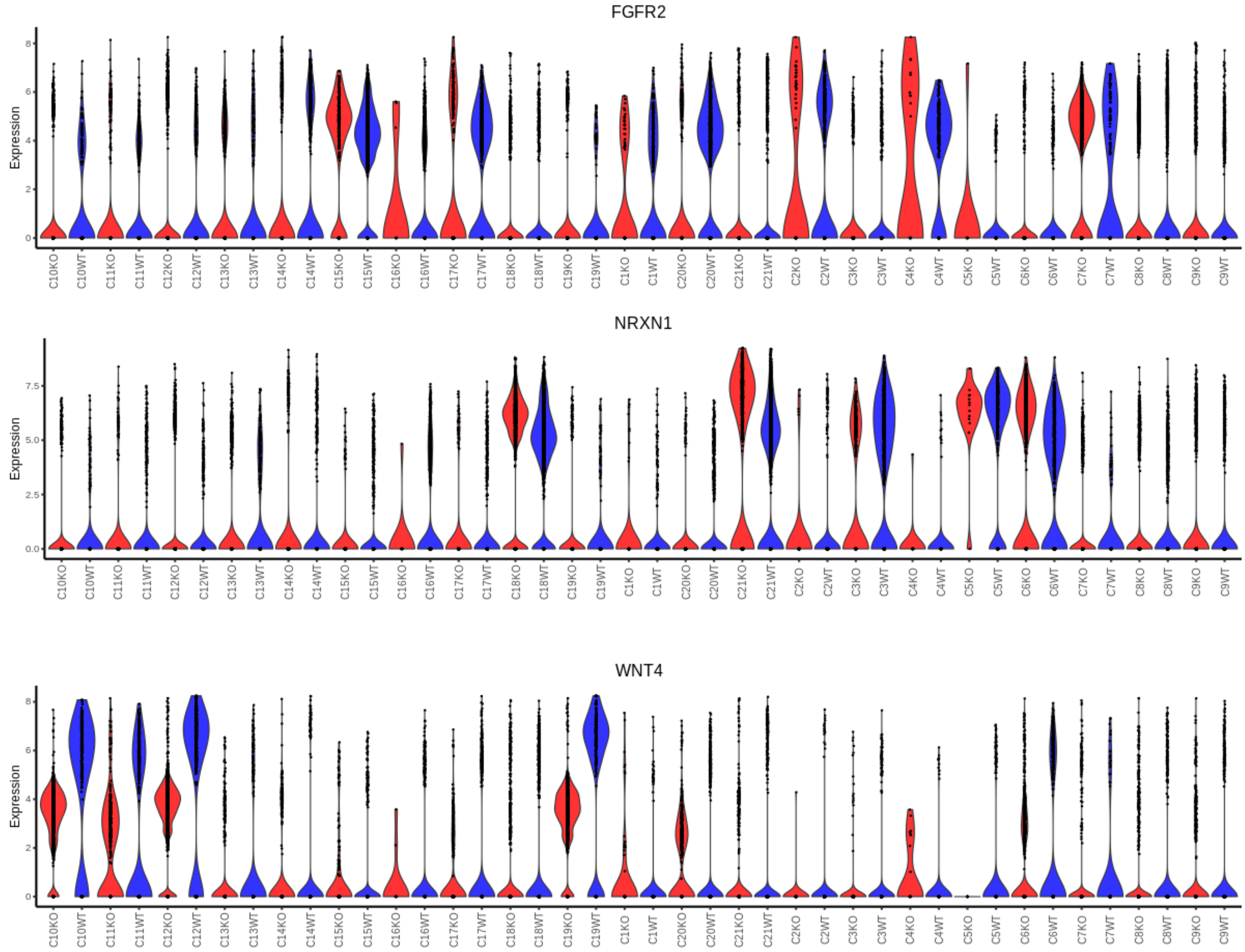


**Red: CHD8 KO cells, Blue: control cells**
